# Prevalence and risk factors of postpartum depression within one year after birth in urban slums of Dhaka, Bangladesh

**DOI:** 10.1101/514729

**Authors:** Rashidul Azad, Rukhshan Fahmi, Sadichhya Shrestha, Hemraj Joshi, Mehedi Hasan, Abdullah Nurus Salam Khan, Mohiuddin Ahsanul Kabir Chowdhury, Shams El Arifeen, Sk Masum Billah

## Abstract

Postpartum depression (PPD) is a serious pubic health concern and known to have the adverse effects on mother’s perinatal wellbeing; an d child’s physical and cognitive development. There were limited literatures on PPD in Bangladesh, especially in urban slum context. The aim of this study was to assess the burden and risk factors of PPD among the urban slum women. A cross-sectional study was conducted between November-December 2017 in three urban slums on 376 women within first 12 months of postpartum. A validated Bangla version of Edinburgh Postnatal Depression Scale was used to measure the depression status. Respondent’s socio-economic characteristics and other risk factors were collected with structured validated questionaire by trained interviewers. Unadjusted Prevalence Ratio (PR) and Adjusted Prevalence Ratio (APR) were estimated with Generalized Linear Model(GLM) and Generalized Estimating Equation (GEE) respectively to identify the risk factors of PPD. The prevalence of PPD was 39.4% within first 12 months following the child birth. Job involvement after child delivery (APR=1.9, 95% CI= 1.1, 3.3), job loss due to pregnancy (APR=1.5, 95% CI= 1.0, 2.1), history of miscarriage or still birth or child death (APR=1.4, 95% CI= 1.0, 2.0), unintended pregnancy (APR=1.8, 95% CI= 1.3, 2.5), management of delivery cost by borrowing, selling or mortgaging assets (APR=1.3, 95% CI= 0.9, 1.9), depressive symptom during pregnancy (APR=2.5, 95% CI= 1.7, 3.8) and intimate partner violence (APR=2.0, 95% CI= 1.2, 3.3), were identified as risk factors. PPD was not associated with poverty, mother in law and any child related factors. The burden of postpartum depression was high in the urban slum of Bangladesh. Maternal mental health services should be integrated with existing maternal health services. Research is required for the innovation of effective, low cost and culturally appropriate PPD case management and preventive intervention in urban slum of Bangladesh.

## Introduction

Postpartum depression (PPD) is a common, non-psychotic mood or mental disorder which typically manifests in mothers within one year of delivery (first year postpartum) [1, 2]. Globally, the prevalence of PPD among mothers ranges from 0.5% to 60.8% [2]. In comparison to women of developed countries, women of developing countries showed higher rates of PPD [3]. A systematic review of 28 developed countries reported that the prevalence PPD symptom (PPDS) was 6-13% among women in high income nations [4]. An independent systematic review on low and middle income countries (LMIC) found the prevalence of postpartum common mental disorder was approximately 20% [5]. Asian countries reported between 3.5-63.3% prevalence rates of depression in postpartum women [6]. In India the prevalence of depression varied from 11% to 16% within fourteen weeks of delivery [7]. Several studies conducted in rural Bangladesh found the prevalence of PPD ranged from 18% to 35% among rural women [8-10].

PPD has negative impacts on children’s physical health. Existing literature of LMICs suggests that PPD leads to poor infant feeding practices, consequently leading to malnutrition and reduced physical growth [11, 12]. The effects of PPD are not limited to physical symptoms but can also impact the mental health of the affected mothers’child. PPD leads to the lower levels of interaction and bonding between the mother and child, which leads to inadequate social, emotional and cognitive development of the child [13, 14]. Research also suggested that repeated level of depression leads to high level of chronic stress at the later age [13]. Further, PPD has a harmful impact on the family and social life [15].

There are various factors causing PPD and the subsequent consequences of PPD varies according to women ages, education levels, races, and ethnicities [16]. Globally, preterm or low birth-weight infants, unemployment, socio-economic deprivation, poor social or emotional support, housing problems, first-born child, child care stress, infant temperament, high parity, obstetric complications, sleep disturbances, low self-esteem, negative attitude towards pregnancy, antenatal depression or anxiety, previous history of depression, poor marital relationship, history of domestic abuse, major adverse life events in preceding one year, neuroticism, perfectionism and level of daily hassles are strong predictors of PPD among women [1, 17-22].

The risk factors identified in the rural areas of Bangladesh were low economic status, nutritional status, physical violence, domestic quarrels with husband and in laws, stress, past mental and depressive symptoms during pregnancy, perinatal death, poor relationship between husband and in laws, morbidity during pregnancy, and current health condition [8, 9, 23, 24]. Comparing rural area, PPD was less explored in urban slum area. We found only one qualitative study explored the cultural attitude towards PPD in urban slum. Of the 36 women interviewed, 17 women were suffering from PPD [25]. In this study women reported that financial crises, adverse life events, intimate partner violence, health problems, and lack of practical social support were causes of daily emotional distress and sadness at postpartum period [25].

The population of urban Dhaka was estimated to be approximately 17 million in 2015 [26] and nearly one third of the population Dhaka city inhabit slums [27]. Within the next 14 years the urban population will increase by 50% due to rapid urbanization [28]. The status of PPD in women in urban slums has received less attention and the prevalence and associated risk factors for PPD in slum areas were unknown. The objective of this proposed study was to assess the burden and risk factors of PPD among women living in urban slums of Dhaka, Bangladesh.

## Materials and methods

### Study design and setting

A cross-sectional study was conducted in three purposively selected large slums from different geographical representative areas in Dhaka city. These three large slums were the Korail slum in Gulshan, the Sattala slum in Mohakhali, and the Mohammadpur slum in Mohammadpur. The reasons for the selection of large slums were; firstly, large slums are typical of other slums in Dhaka city in terms of household structure, overall environment, religion and culture; secondly, the population of large slum is heterogeneous in terms of place of origin; thirdly, the area chosen was large enough to get sufficient research participants for the study [29]. In addition, three different slums from three different areas addressed the lack of variation that could arise from selecting the slums from only one area.

### Study participants, sample size and sampling strategy

The study population were postpartum mothers with a child under the age of one year living in urban slums. Considering 22% PPD prevalence in rural area of Bangladesh [8]; and with 4.2% margin of error and 95% level of significance, the calculated sample size was 374. We divided each slum into different sectors and physically visited all households of each sector and searched the postpartum mothers according to our inclusion criteria. If mothers were found, present and agreed to participate in our study then we interviewed those mothers. One local woman assisted the data collectors in the door to door searching of postpartum mothers. We visited and searched for study participants in all household of Sattala and Koril slum; and partially some sectors of Mohammadpur slum; and ended up with 376 interviews of postpartum mothers.

### Data collection

The data we collected between November and December 2017. The data collection team included four research team members and four research assistants with backgrounds in social sciences. Experienced research psychologist, gender experts and maternal and child health (MCH) experts intensively trained the data collection team. Each day after data collection a team meeting was held and filled questionnaires were cross-checked. Possible erratic responses were identified, discussed and corrected.

### Outcome measure

The Edinburgh Postnatal Depression Scale (EPDS) was used to detect depressive symptoms during the postpartum period [30]. EPDS is a 10-item questionnaire assessing the interviewee’s depressive feelings from the seven days before the interview. The response of the questions is scored from 0-3, with higher scores indicating higher levels of depressive symptoms. The total EPDS score of a respondentcan range from 0 to 30. Cox et al. (1987) proposed a cutoff level ten (≥10) to reduce the detection failure of PPD. In Bangladesh, a validation study showed 89% sensitivity, 87% specificity, 40% positive predictive value and 99% negative predictive value of using 10 as the cutoff score [31]. Another follow-up study of EPDS usage in Bangladesh demonstrated good reliability with Cronbach’s alpha 0.70 and 0.75 [9]. In this study, greater than the 10 cutoff value of EPDS scale was used to screen PPD, which was consistent with other studies in Bangladesh [8-10, 23].

### Risk factors

The validated tools of Bangladesh Demographic and Health Survey (BDHS), Bangladesh Urban Health Survey (BUHS) and WHO Domestic Violence Against Women were adapted to prepare our tools of the study. The data were collected in the following manner for the analysis of risk factors.

#### Socio-economic characteristics

Socio-demographic information (age, age of marriage, educational status, income generating activities before pregnancy and after pregnancy, etc) and household characteristics (basic household construction, ownership, availability of toilet and water facilities, household belongings, etc) were collected through face-to-face interviews using the standard validated questionnaire of BUHS [28]. The wealth score of households were computed through principal component analysis using household characteristics of respondents. During the calculation of wealth score, we included all poverty related indicators of household, which included the ownership of household, household construction materials, type and sharing status of toilet facility and water source, and household assets.

#### Pregnancy related factors

Obstetric, reproductive, and child characteristics such as the number of children, history of miscarriage and child death, pregnancy intention, age and sex of last child, reported gestational age and child birth weight, neo-natal complications, ANC and PNC information, mode of delivery (caesarean or normal), cost of delivery, and delivery cost management were collected through standard questions. Gestational ages less than 37 weeks were categorized as pre-term birth and birth weight less than 2.5 kg was considered to be low birth weight.

Each respondent was asked whether she had experienced at least one depressive symptom of EPDS scale during pregnancy period. If their response was positive then we categorized them as depressive symptom started to develop from pregnancy period. We also asked respondents whether family, work or any other form mental stress took place during pregnancy period. This was characterized as perceived antenatal stress.

#### Family support and intimate partner violence

Women’s experience of intimate partner violence (IPV) was collected using the domestic violence module of WHO’s multicounty study, which included Bangladesh as well [32]. The study divided intimate partner violence into three categories; psychological, physical and sexual. Each respondent was asked if they any type of IPV during their last pregnancy and 12 months before the conception.

Family support was measured by assistance level of mother in law and husband in taking care of children and helping in household chores. This was assessed through likert scale (always, often, sometime, rarely and never). Relationship with husband was assessed by asking the sharing status of personal feelings to husband with similar likert scale.

### Data analysis

We calculated the descriptive statistics of respondents’ background characteristics, PPD status, and other relevant variables. Then we conducted a series of cross-tabulation including the chi-square test to check the bivariate relationship of PPD and relevant risk factors (p-value of chi square test was not reported in the data table). Prevalence Ratio (PR) was used as a measure of relative strength of association of different risk factor of this cross sectional study [33-35]. PR of bivariate analysis was calculated using Generalized Linear Model (GLM) with poisson family and log link.

Risk factors identified in the bivariate analysis were analyzed in the multivariable model. Risk factors were added to the multivariate model according to the strength of association in bivariate analysis. Moreover, multi-collinearly and overlapping nature among the variables were considered for selection of risk factors in the multivariate model. Adjusted analysis (clustering and confounder adjustment) was conducted using the Generalized Estimating Equation (GEE). Statistical software STATA version 13 was used in data analysis.

### Ethical consideration

Prior to the study, ethical approval was taken from the Ethical Review Committee (ERC), JPGSPH. Participants were recruited after reading the full statement of consent and signing the Bangla written consent form. In urban slum, most of the family is single family and parent/guardian was not always available in house due to their job engagement. For that reason, with the prior knowledge and approval of ERC, we had not any provision of parental or guardian consent for the mothers under 18 year age. However, if guardian was present in house then we took his/her verbal consent. We referred the depressed mothers or who had the thought of harming herself to the Government’s National Mental Institute of Bangladesh.

## Results

Among the post-partum mothers 25.2% were adolescents and 31.7% were illiterate or with just enough skills to sign their names. On the other hand, 17.0% of mothers attained a secondary level of education or received a higher education. Approximately half, (48.7%) were first time mothers and 20.0% had three or more children. The mean number of household members was 4.5 and most of the families (83.8%) resided in one room. Only 7.7% of mothers were working after delivering their child, but before their most recent pregnancy around 49.7% of mothers had been engaged in income generating activities. Among the respondents employed before pregnancy, 54.6% were garments or industry workers, 27.3% were part-time house maid; whereas, 10.7% worked in private and other non-government organization and 7.0% involved some sort of home based work (Tailoring, handicraft, etc). Among the respondents we interviewed, 22.6% experienced at least one child death, or miscarriage, or stillbirth during her entire fertility period and 67.6% of respondents planned their most recent pregnancy. About 69.1% of mothers faced intimate partner violence before their last pregnancy, and 47.7% faced it during the pregnancy period as well (Table 1).

**Table 1:** Respondent’s characteristics

| Indicators | n(%), Mean±SD<br>(N=376) |
| --- | --- |
| Respondent's age |  |
| 13-19 | 85 (25.2) |
| 20-29 | 208 (61.7) |
| 30-49 | 44 (13.1) |
| Education of respondent |  |
| No education/signed only | 119 (31.7) |
| Primary | 193 (51.3) |
| Secondary or higher | 64 (17.0) |
| No of children |  |
| 1 | 183 (48.7) |
| 2 | 118 (31.4) |
| 3+ | 75 (20.0) |
| Mean number of household members | 4.5± 0.09 |
| No of living room |  |
| 1 | 315 (83.8) |
| 2+ | 61 (16.2) |
| Job involvement before pregnancy | 187 (49.7) |
| Type of job involved before pregnancy |  |
| Private/NGO job | 20 (10.7) |
| Public job | 1 (0.5) |
| Garments/industry worker | 102 (54.6) |
| Part-time house maid | 51 (27.3) |
| Tailoring, handicraft or other home based work | 13 (7.0) |
| Currently working | 29 (7.7) |
| Job loss due to pregnancy | 164 (47.3) |
| History of miscarriage or still birth or child death | 85 (22.6) |
| Age of the last child |  |
| <3 months | 111 (29.6) |
| 3<6 months | 113 (30.1) |
| 6<9 months | 79 (21.1) |
| 9<12 months | 72 (19.2) |
| Intended pregnancy | 254 (67.6) |
| Gestational age |  |
| Pre-term birth | 58 (15.8) |
| Term birth | 309 (84.2) |
| Birth weight of last child |  |
| Low birth weight | 161 (47.9) |
| Normal weight | 175 (52.1) |
| Neo-natal complications | 78 (21.7) |
| Cost of child delivery managed by borrowing/sold asset/mortgage | 96 (25.5) |
| Intimate partner violence before last pregnancy* | 259 (69.1) |
| Intimate partner violence during last pregnancy* | 179 (47.7) |
| Postpartum depression status | 148 (39.4) |
| At least one depressive symptom reported to develop during pregnancy | 189 (50.3) |
| Perceived antenatal stress | 143 (38.0) |

\*At least one type of psychological, physical or sexual violence

### Prevalence of postpartum depression

After administration of EPDS scale, 148 (39.4%) mothers out of 376 mothers were found having postpartum depression. About 50.3% mothers reported at least one symptom of EPDS scale developed during pregnancy period and 38.0% reported any family, working and other form of mental stress during the pregnancy period (Table 1).

### Potential risk factors of postpartum depression

The potential risk factors of postpartum depression were organized in following categories: socio-economic factors; pregnancy related factors and family support and intimate partner violence (Table 2).

**Table 2:** Bivariate and multivariable association of postpartum depression with socio-economic factors, pregnancy related factors, and family support and intimate partner violence

| Indicators | n | Depressed (%) | Prevalence Ratio (PR) (95% CI) | p-value | Adjusted Prevalence Ratio (APR)<sup>‡</sup>(95% CI) | p-value |
| --- | --- | --- | --- | --- | --- | --- |
| **Socio-economic factors** |  |  |  |  |  |  |
| Respondent's age |  |  |  |  |  |  |
| 13-19 | 85 | 32.9 | 1 |  |  |  |
| 20-29 | 208 | 42.3 | 1.3 (0.8,2.0) | 0.249 |  |  |
| 30-49 | 44 | 54.6 | 1.7 (0.9,2.6) | 0.070 |  |  |
| Wealth quintile |  |  |  |  |  |  |
| 1st quintile<br>(Comparatively poor) | 76 | 40.8 | 1.2 (0.7,<br>2.1) | 0.453 |  |  |
| 2nd quintile | 75 | 49.3 | 1.5 (0.9,<br>2.5) | 0.130 |  |  |
| 3rd quintile | 75 | 34.7 | 1.0 (0.6,<br>1.8) | 0.889 |  |  |
| 4th quintile | 75 | 38.7 | 1.2 (0.7,<br>2.0) | 0.587 |  |  |
| 5th quintile<br>(Comparatively rich) | 75 | 33.3 | 1 |  |  |  |
| Education of respondent |  |  |  |  |  |  |
| No education/signed<br>only | 119 | 48.7 | 1.8 (1.1,3.2) | 0.028 | 1.2 (0.7,2.1 ) | 0.559 |
| Primary | 193 | 37.8 | 1.4 (0.8,2.4) | 0.189 | 0.9 (0.5,1.6 ) | 0.687 |
| Secondary or higher | 64 | 26.6 | 1 |  | 1 |  |
| Current profession |  |  |  |  |  |  |
| Working | 29 | 62.1 | 1.7 (1.0,2.7) | 0.045 | 1.9 (1.1,3.3 ) | 0.020 |
| Not working | 347 | 37.5 | 1 |  | 1 |  |
| Job loss due to pregnancy |  |  |  |  |  |  |
| Yes | 164 | 45.7 | 1.5 (1.1,2.2) | 0.018 | 1.5 (1.0,2.1 ) | 0.040 |
| No | 183 | 30.1 | 1 |  | 1 |  |
| No of children |  |  |  |  |  |  |
| 1 | 183 | 33.9 | 1 |  | 1 |  |
| 2 | 118 | 39.8 | 1.2 (0.8,1.7) | 0.403 | 1.0 (0.7,1.5 ) | 0.982 |
| 3+ | 75 | 52.0 | 1.5 (1.0,2.3) | 0.036 | 1.0 (0.6,1.5 ) | 0.843 |
| **Pregnancy related factors** |  |  |  |  |  |  |
| History of miscarriage or<br>still birth or child death |  |  |  |  |  |  |
| Yes | 85 | 50.6 | 1.4 (1.0,2.0) | 0.062 | 1.4 (1.0,2.0 ) | 0.073 |
| No | 291 | 36.1 | 1 |  | 1 |  |
| Pregnancy type |  |  |  |  |  |  |
| Intended pregnancy | 254 | 29.9 | 1 |  | 1 |  |
| Unintended pregnancy | 122 | 59.0 | 2.0 (1.4,2.7) | <0.001 | 1.8 (1.3,2.5 ) | 0.001 |
| Gestational age |  |  |  |  |  |  |
| Pre-term birth | 58 | 48.3 | 1.3 (0.8,1.9) | 0.265 |  |  |
| Term birth | 309 | 38.2 | 1 |  |  |  |
| Birth weight of last child |  |  |  |  |  |  |
| Low birth weight | 161 | 42.2 | 1.1 (0.8,1.5) | 0.627 |  |  |
| Normal weight | 175 | 38.9 | 1 |  |  |  |
| Age of the last child |  |  |  |  |  |  |
| <3 months | 111 | 38.7 | 1 |  |  |  |
| 3<6 months | 113 | 35.4 | 0.9 (0.6,1.4) | 0.681 |  |  |
| 6<9 months | 79 | 43.0 | 1.1 (0.7,1.7) | 0.647 |  |  |
| 9<12 months | 72 | 43.1 | 1.1 (0.7,1.8) | 0.654 |  |  |
| Neo-natal complications |  |  |  |  |  |  |
| Yes | 78 | 46.2 | 1.2 (0.9,1.8) | 0.254 |  |  |
| No | 281 | 37.0 | 1 |  |  |  |
| Child delivery cost management by borrowing/sold asset/mortgage |  |  |  |  |  |  |
| Yes | 96 | 55.2 | 1.6 (1.2,2.3) | 0.005 | 1.3 (0.9,1.9 ) | 0.108 |
| No | 280 | 33.9 | 1 |  | 1 |  |
| Perceived antenatal stress |  |  |  |  |  |  |
| Yes | 143 | 65.7 | 2.8 (2.0,4.0) | <0.001 |  |  |
| No | 233 | 23.2 | 1 |  |  |  |
| At least one depressive symptom reported to develop during pregnancy |  |  |  |  |  |  |
| Yes | 189 | 61.4 | 3.6 (2.4,5.3) | <0.001 | 2.5 (1.7,3.8 ) | <0.001 |
| No | 187 | 17.1 | 1 |  | 1 |  |
| **Family support and intimate partner violence** |  |  |  |  |  |  |
| Mother-in-law take care of child and help in household work |  |  |  |  |  |  |
| Always/Often | 64 | 40.6 | 1 |  |  |  |
| Sometimes/Rarely/ Never | 17 | 64.7 | 1.6 (0.8,3.2) | 0.196 |  |  |
| Husband take care of child and help in household work |  |  |  |  |  |  |
| Always/Often | 230 | 36.5 | 1 |  |  |  |
| Sometimes /Rarely/Never | 108 | 50.0 | 1.4 (1.0,1.9) | 0.072 |  |  |
| Share personal feelings with husband |  |  |  |  |  |  |
| Always/Often | 271 | 33.2 | 1 |  |  |  |
| Sometimes/ Rarely/ Never | 67 | 71.6 | 2.2 (1.5,3.1) | <0.001 |  |  |
| Intimate partner violence before last pregnancy |  |  |  |  |  |  |
| Yes | 259 | 49.4 | 3.0 (1.9,4.9) | <0.001 |  |  |
| No | 116 | 16.4 | 1 |  |  |  |
| Intimate partner violence during last pregnancy |  |  |  |  |  |  |
| Yes | 179 | 55.3 | 2.3 (1.6,3.2) | <0.001 |  |  |
| No | 196 | 24.5 | 1 |  |  |  |
| Intimate partner violence before or during pregnancy period |  |  |  |  |  |  |
| Yes | 260 | 49.6 | 3.0 (1.9,4.9) | <0.001 | 2.0 (1.2,3.3 ) | 0.009 |
| No | 116 | 16.4 | 1 |  | 1 |  |

¥Estimates adjusted for clustering at slum level and wealth score

### Bivariate risk factor analysis

#### Socio-economic factors

Postpartum depression among the women of urban slum was not associated with respondent age and wealth quintile (p>0.050). The prevalence of postpartum depression was higher in the uneducated group of women (48.7%) compare to the women (26.6%) who completed the secondary or higher level of education (PR=1.8, p=0.028, 95% CI= 1.1, 3.2). The prevalence of depression was more common in currently employed mothers comparing unemployed mothers (PR=1.7, p=0.045, 95% CI= 1.0, 2.7). The likelihood of depression among the mothers who worked before pregnancy but left their work due to pregnancy is also higher than the likelihood of depression in other groups of mothers (PR=1.5, p=0.018, 95% CI= 1.1, 2.2).

#### Pregnancy related factors

Respondents who experienced miscarriage or still birth or child death had a higher prevalence of postpartum depression (PR=1.4, p=0.062, 95% CI= 1.0, 2.0). Unplanned pregnancy was a major factor of postpartum depression among the study participants (PR=2.0, p<0.001, 95% CI= 1.4, 2.7). Postpartum depression were not correlated with factors related to the child such as preterm birth (p=0.265), low birth weight child (p=0.627) and neonatal complication (p =0.254). However, the prevalence of depression was 60% higher for mothers, whose family arranged the delivery cost by borrowing, selling or through mortgage assets (p=0.005). Perceived antenatal stress (family, working or any other mental stress) during pregnancy was another strong predictor of PPD (PR=2.8, p<0.001, 95% CI= 2.0, 4.0). PPD was most common among the mothers who had at least one EPDS depressive symptom developed during pregnancy period (PR=3.6, p<0.001, 95% CI= 2.4, 5.3).

#### Family support and intimate partner violence

Maternal depression was not associated with mother-in-law’s support in slum area (p=0.196), but depression was fairly associated with the practical support of husband (p=0.072). The prevalence of depression was more than two times higher for women who sometimes/never/rarely shared their personal feelings with their husband in comparison to the women who always or often shared their personal feelings with their husband (PR=2.2, p<0.001, 95% CI= 2.0, 4.5). Similarly, PPD was most common for women who faced the intimate partner violence both before their recent pregnancy and during the pregnancy period [(IPV before pregnancy: PR=3.0, p<0.001, 95% CI= 1.9, 4.9); (IPV during pregnancy: PR=2.3, p<0.001, 95% CI= 1.6, 3.2)].

### Multivariable risk factor analysis

After inclusion of all relevant factors in multivariate regression model, the risk estimates slightly reduced but remained significant for current job involvement, job loss due to pregnancy, unintended pregnancy, development of depressive symptoms during pregnancy period, and intimate partner violence. In the full model, the adjusted prevalence of depression was almost 2 times higher (APR=1.9, p=0.020, 95% CI=1.1, 3.3) for those who had some form of employment and 50% higher for those who quit their job due to pregnancy (APR=1.5, p=0.040, 95% CI=1.0, 2.1). For the postpartum mothers who conceived unintentionally, the adjusted prevalence of depression was 80% higher (APR=1.8, p=0.001, 95% CI= 1.3, 2.5). The adjusted prevalence of postpartum depression was about 2.5 times higher among the women who reported to develop the depressive symptom during the pregnancy period (APR=2.5, p<0.001, 95% CI=1.7, 3.8). The adjusted prevalence of depression was 2 times higher (APR=2.0, p=0.009, 95% CI= 1.2, 3.3) among the postpartum mothers who faced intimate partner violence before child birth.

However, other risk factors, i.e., delivery cost management by borrowing/selling asset/mortgage (APR=1.3, 95% CI=1.0, 1.9) and history of miscarriage or still birth or child death (APR=1.4, 95% CI=1.0, 2.0) were closely significant at 5% level of significance, which may be significant for larger samples of postpartum mothers in urban slum.

## Discussion

This study explored the burden of postpartum depression and the associated factors in slum areas. Our study results show that about 40 women out of 100 women were suffering from PPD and the associated risk factors were current job involvement, job loss due to pregnancy, history of miscarriage, still birth and child death, unintended pregnancy, cost of delivery managed from borrowing/selling asset/mortgage, depressive symptom during pregnancy period, perceived antenatal stress, poor marital relationship with husband, and intimate partner violence.

The burden of PPD in women who inhabited slums was higher in comparison that of postpartum mothers living in rural areas [8-10]. Existing mental disorder data of urban slum women also suggested the higher level of mental disease burden in slum area. In one recent study at an urban slum, Khan et al. found that 46% mothers with children under the age of five suffered from common mental disorders [36].

Half of our respondents had been involved in income generating activities before the pregnancy, while only eight percent of survey respondents were involved in income generating activities during the postpartum period. Depression was more common for the women who worked but quit job due to pregnancy. There was the scarcity of data about the postpartum mental health and unemployment in low income countries. In a study of Canadian postpartum mothers, depression was less common in women who were in maternity leave [37]. Similarly, an Australian study found the evidence that paid maternal leave was beneficial for post-partum health and wellbeing [38]. According to the Labor Force Survey (2016-17), 89.1% female employment in urban Dhaka was informal employment [39]. In Bangladesh Labour Law there is a provision for paid maternity leave [40]. A strict implementation of clause 45 to 50 of the Bangladesh Labour Law might improve the postpartum mental health of slum working women. We found that more than half of our respondents worked in garments or industry sector, however, the status of paid maternity leave was not satisfactory in garments industry [41, 42]. Among the informal employment garments and other industries were easy to track. So, at the first stage, at least in the garments and industry sector, the strict enforcement of paid maternity leave needed to initiate to reduce the burden of PPD in slum.

Postpartum depression was also high among the mothers who were working after child delivery. This was because with the professional working stress, taking care of new child added additional stress to women. These multiple roles leaded to role overload that could have negative effects on psychological well-being of postpartum mother [37, 43]. Some government oversight of maternity schemes for postpartum women, especially for the women working in slum areas, might improve the postpartum mental wellbeing of slum women.

PPD was not associated with household economic status within slum areas. This result contradicts the PPD findings within the rural context of Bangladesh [9, 24]. The possible explanation could be the people of slum area were concerned with immediate economic survival [44] rather than long term economic stability (measured by the wealth index).

However, debt and sudden financial hardship of family may affect the postpartum mental health of women [45, 46]. Depression was higher for women, whose child delivery cost was managed by borrowing, selling assets or through mortgages. The findings of recent national maternal mortality survey revealed that private sector delivery has increased several fold and has the highest delivery cost [47]. To minimize the costs of delivery, necessary steps are needed to promote deliveries in public and NGOs facilities.

In rural area of Bangladesh, association of postpartum depression with unintended pregnancy was found in bivariate analysis, but the association disappeared in multivariate regression analysis [8, 9, 23]. However, in the urban slum context, the unintended pregnancy was an independent risk factor for PPD. The association of PPD with unintended pregnancy persisted even after controlling other risk factors in the multivariate model. An unmet need of contraceptives was found in both urban and rural areas [48]. A patient centred approach and providing comprehensive information and access to contraceptive options may helpful for controlling the unintended pregnancy [49].

Among the obstetric factors assessed in our study, a history of miscarriage or still birth or child death was associated with postpartum depression; a finding that was consistent with those of other studies [8, 50, 51]. The fear of a repeat occurrence of a miscarriage, stillbirth, or child death are thought to be a contributory factors of depression in the antenatal, postnatal period, or in both periods. The recommended number of antenatal care (ANC) and postnatal care (PNC) visits may reduce miscarriage, still birth and postnatal child death; and consequently reduce PPD rates in subsequent pregnancies. Healthcare providers should consider the history of earlier loss as a factor of increased vulnerability of depression during the next pregnancy and early postpartum period [52].

Depressive symptom during pregnancy was mostly associated with postpartum depression, similar to the findings of other studies [4, 8, 53]. Perceived family, working, and any other mental stress during pregnancy was also associated with PPD like other findings [54, 55]. Perceived antenatal stress may contribute to development of depressive symptoms in the antenatal period [56].These findings highlighted PPD is not only a matter of postpartum period and noted the importance of intervention in the prenatal period to prevent PPD [53, 54].

Intimate partner violence (IPV) is the one of the most important risk factors found in our study, which was consistent with most of the literatures of postpartum depression [9, 23, 57-62]. In a recent study of Bangladesh, Ziaie found that all forms of domestic violence were strongly associated with higher levels of emotional distress during the pregnancy period as well [63]. In our study, we also found that poor marital relationship (sharing personal feelings and practical support in household work) with husbands was a strong predictor of PPD, consistent with the findings of other studies [8, 22]. IPV and poor marital relationship might be associated with each other [64]. Therefore, these findings suggested that couples focused intervention from the prenatal period may reduce the risk of PPD.

## Strengths and limitations

The main strength of the study was the study focused the rapid increasing segment of urban population of Bangladesh with a large sample size. To our knowledge it was the first study to assess the prevalence and risk factors of postpartum depression in urban slum of Bangladesh. In addition, the study included the wide range of risk factors found in the postpartum literatures of Bangladesh and similar context. We used GLM and GEE model to estimate Prevalence Ratio (PR) and to adjust the clustering effect at slum level. Statistically, estimating PR has some methodological advantages as it is a comparable measure of relative risk (RR) for cross-sectional data, and the estimation models of PR able to control the confounders and interactions more adequately than the Logistic regression models [65, 66]. Most importantly, the study used the repeatedly validated Bangla version of instruments and scale which was nationally and internationally recognized and widely used.

The study had some limitations worth to acknowledge. The design of the study was cross-sectional and therefore could not measure the incidence of PPD. Data about most of the risk factors were collected from mother’s recall, which may lead to under or over reporting of symptoms. However, data collectors were intensively trained on the interview techniques to reduce the recall bias. In Dhaka city, there were about 4,966 slums of different sizes [67]. In this study, we purposively selected 3 large slums, as slum level randomization was not possible due to time and resource constraint. This may lead some selection bias in terms of representativeness of slums of Dhaka city. However, in terms of generalisability of study results, this selection bias was less, as overall contextual factors of slums are representatively prevailing in the large slums of Dhaka city. In addition, the frequent movement of slum dwellers across the slums of [28, 68] keeps the urban slums typically heterogeneous. We used a likert scale to measure the support of mother in law and husband to the mother, which were not validated before our use. However, we believe that this likert scale was a good instrument to measure the ‘Family Support’. Some working mothers may not be reached due to strict data collection timeline and not having schedule of household revisit. We tried to overcome this constraint by extending the data collection time beyond the office hour and collecting data in weekend as well.

## Conclusions

The higher prevalence of PPD suggested the importance of mental health support system for the low income women in slum area. Maternal mental health services should be integrated with existing maternal health services. The primary maternal health care staffs could be provided the basic PPD screening and its primary management training, so that they can refer the PPD cases for appropriate mental health services when needed. They are also needed to educate about the contextually relevant risk factors of PPD as part of the training component. Additionally, the existing maternal health services in slum area should be strengthened and pro-poor friendly.

Moreover, research is required to develop the low cost non-pharmacological management of PPD cases in the informal settlement of urban poor. In addition to that, research for developing the culturally appropriate preventive interventions to control the risk factors should be undertaken in the urban slum of Bangladesh.

## Acknowledgements

The study was supported by the James P. Grant School of Public Health (JPGSPH). This paper is based on the data collected for Summative Learning Project (SLP) for author’s partial requirement of Master of Public Health (MPH) degree of JPGSPH at BRAC University. The author is grateful to Mahfuz Al Mamun, gender expert and Farzana Ramee, research psychologist of icddr,b for their training support. He is also thankful to Aniqa Hassan for her final language editing of author’s manuscript. Finally he wants to thank icddr,b for providing the scholarship opportunity for pursuing the MPH degree in JPGSPH.

## Supporting information

**S1 File. Data file.**

(DTA)

**S2 File. Regression Model Parameters.**

(DOCX)

## References

1. Stewart DE, Robertson E, Dennis C-L, Grace SL, Wallington T. Postpartum depression: Literature review of risk factors and interventions. Toronto: University Health Network Women’s Health Program for Toronto Public Health. 2003.

2. Halbreich U, Karkun S. Cross-cultural and social diversity of prevalence of postpartum depression and depressive symptoms. Journal of affective disorders. 2006;91(2):97–111.

3. Tronick E, Reck C. Infants of depressed mothers. Harvard review of psychiatry. 2009;17(2):147–56.

4. Gavin NI, Gaynes BN, Lohr KN, Meltzer-Brody S, Gartlehner G, Swinson T. Perinatal depression: a systematic review of prevalence and incidence. Obstetrics & Gynecology. 2005;106(5, Part 1):1071–83.

5. Fisher J, Mello MCd, Patel V, Rahman A, Tran T, Holton S, et al. Prevalence and determinants of common perinatal mental disorders in women in low-and lower-middle-income countries: a systematic review. Bulletin of the World Health Organization. 2012;90(2):139–49.

6. Klainin P, Arthur DG. Postpartum depression in Asian cultures: a literature review. International journal of nursing studies. 2009;46(10):1355–73.

7. Hegde S, Latha K, Bhat SM, Sharma P, Kamath A, Shetty A. Postpartum depression: prevalence and associated factors among women in India. J Womens Health Issues Care. 2012;1(1):1–7.

8. Gausia K, Fisher C, Ali M, Oosthuizen J. Magnitude and contributory factors of postnatal depression: a community-based cohort study from a rural subdistrict of Bangladesh. Psychological medicine. 2009;39(6):999–1007.

9. Nasreen HE, Edhborg M, Petzold M, Forsell Y, Kabir ZN. Incidence and risk factor of postpartum depressive symptoms in women: a population based prospective cohort study in a rural district in Bangladesh. J Depress Anxiety. 2015;4(1000180):2167–1044.1000180.

10. Islam MJ, Baird K, Mazerolle P, Broidy L. Exploring the influence of psychosocial factors on exclusive breastfeeding in Bangladesh. Archives of women’s mental health. 2017;20(1):173–88.

11. Rahman A, Harrington R, Bunn J. Can maternal depression increase infant risk of illness and growth impairment in developing countries? Child: care, health and development. 2002;28(1):51–6.

12. Anoop S, Saravanan B, Joseph A, Cherian A, Jacob K. Maternal depression and low maternal intelligence as risk factors for malnutrition in children: a community based case-control study from South India. Archives of Disease in Childhood. 2004;89(4):325–9.

13. Hammen C. Social stress and women’s risk for recurrent depression. Archives of Women’s Mental Health. 2003;6(1):9–13.

14. Murray L, Cooper PJ. Postpartum depression and child development. Psychological medicine. 1997;27(2):253–60.

15. O’hara MW, Swain AM. Rates and risk of postpartum depression—a meta-analysis. International review of psychiatry. 1996;8(1):37–54.

16. Dennis CL, Chung-Lee L. Postpartum depression help-seeking barriers and maternal treatment preferences: A qualitative systematic review. Birth. 2006;33(4):323–31.

17. Vigod SN, Villegas L, Dennis CL, Ross LE. Prevalence and risk factors for postpartum depression among women with preterm and low-birth-weight infants: a systematic review. BJOG: An International Journal of Obstetrics & Gynaecology. 2010;117(5):540–50.

18. Milgrom J, Gemmill AW, Bilszta JL, Hayes B, Barnett B, Brooks J, et al. Antenatal risk factors for postnatal depression: a large prospective study. Journal of affective disorders. 2008;108(1):147–57.

19. Robertson E, Grace S, Wallington T, Stewart DE. Antenatal risk factors for postpartum depression: a synthesis of recent literature. General hospital psychiatry. 2004;26(4):289–95.

20. Norhayati M, Hazlina NN, Asrenee A, Emilin WW. Magnitude and risk factors for postpartum symptoms: a literature review. Journal of affective Disorders. 2015;175:34–52.

21. Nielsen D, Videbech P, Hedegaard M, Dalby J, Secher NJ. Postpartum depression: identification of women at risk. BJOG: An International Journal of Obstetrics & Gynaecology. 2000;107(10):1210–7.

22. Beck CT. Predictors of postpartum depression: an update. Nursing research. 2001;50(5):275–85.

23. Islam MJ, Broidy L, Baird K, Mazerolle P. Intimate partner violence around the time of pregnancy and postpartum depression: The experience of women of Bangladesh. PloS one. 2017;12(5):e0176211.

24. Surkan PJ, Sakyi KS, Christian P, Mehra S, Labrique A, Ali H, et al. Risk of Depressive Symptoms Associated with Morbidity in Postpartum Women in Rural Bangladesh. Maternal and child health journal. 2017;21(10):1890–900.

25. Williams A, Sarker M, Ferdous ST. Cultural Attitudes toward Postpartum Depression in Dhaka, Bangladesh. Medical anthropology. 2018;37(3):194–205.

26. Khalequzzaman M, Chiang C, Hoque BA, Choudhury SR, Nizam S, Yatsuya H, et al. Population profile and residential environment of an urban poor community in Dhaka, Bangladesh. Environmental health and preventive medicine. 2017;22(1):1.

27. Perry HB. Health for all in Bangladesh: lessons in primary health care for the twenty-first century: University Press; 2000.

28. National Institute of Population Research and Training (NIPORT), International Centre for Diarrhoeal Disease Research, Bangladesh (icddr,b) and MEASURE Evaluation. Bangladesh Urban Health Survey 2013 Final Report. Dhaka, Bangladesh and Chapel Hill, North Carolina, USA: NIPORT, icddr,b, and MEASURE Evaluation; 2015.

29. Uzma A, Underwood P, Atkinson D, Thackrah R. Postpartum health in a Dhaka slum. Social Science & Medicine. 1999;48(3):313–20.

30. Cox JL, Holden JM, Sagovsky R. Detection of postnatal depression: development of the 10-item Edinburgh Postnatal Depression Scale. The British journal of psychiatry. 1987;150(6):782–6.

31. Gausia K, Fisher C, Algin S, Oosthuizen J. Validation of the Bangla version of the Edinburgh Postnatal Depression Scale for a Bangladeshi sample. Journal of Reproductive and Infant Psychology. 2007;25(4):308–15.

32. García-Moreno C, Jansen HAFM, Ellsberg M, Heise L, Watts C. WHO multi-country study on women’s health and domestic violence against women. World Health Organization (WHO), 2005.

33. Barros AJ, Hirakata VN. Alternatives for logistic regression in cross-sectional studies: an empirical comparison of models that directly estimate the prevalence ratio. BMC medical research methodology. 2003;3(1):21.

34. Deddens J, Petersen M. Approaches for estimating prevalence ratios. Occupational and environmental medicine. 2008;65(7):501–6.

35. skove T, Deddens J, Petersen MR, Endahl L. Prevalence proportion ratios: estimation and hypothesis testing. International journal of epidemiology. 1998;27(1):91–5.

36. Khan AM, Flora MS. Maternal common mental disorders and associated factors: a cross-sectional study in an urban slum area of Dhaka, Bangladesh. International journal of mental health systems. 2017;11(1):23.

37. des Rivières-Pigeon C, Séguin L, Goulet L, Descarries F. Unravelling the complexities of the relationship between employment status and postpartum depressive symptomatology. Women & Health. 2001;34(2):61–79.

38. Hewitt B, Strazdins L, Martin B. The benefits of paid maternity leave for mothers’ post-partum health and wellbeing: Evidence from an Australian evaluation. Social Science & Medicine. 2017;182:97–105.

39. Bangladesh Bureau of Statistics. Report on Labour Force Survey (LFS) 2016-17. Report. Bangladesh: Bangladesh Bureau of Statistics, 2018.

40. Ministry of Law. Bangladesh Labour Law, 2006. Legislative and Parliamentary Affairs Division, Justice and Parliamentary Affairs, Ministry of Law, Bangladesh; 2006.

41. Akhter S, Salahuddin A, Iqbal M, Malek A, Jahan N. Health and occupational safety for female workforce of garment industries in Bangladesh. Journal of Mechanical Engineering. 2010;41(1):65–70.

42. Bearnot E. Bangladesh: A labor paradox. World Policy Journal. 2013;30(3):88–97.

43. Haggag AK, Geser W, Ostermann H, Schusterschitz C. Depressive symptoms in mothers: The role of employment and role quality. Journal of Workplace Behavioral Health. 2011;26(4):313–33.

44. Islam MM, Ali M, Ferroni P, Underwood P, Alam MF. Prevalence of psychiatric disorders in an urban community in Bangladesh. General hospital psychiatry. 2003;25(5):353–7.

45. Reading R, Reynolds S. Debt, social disadvantage and maternal depression. Social science & medicine. 2001;53(4):441–53.

46. Braveman P, Marchi K, Egerter S, Kim S, Metzler M, Stancil T, et al. Poverty, near-poverty, and hardship around the time of pregnancy. Maternal and Child Health Journal. 2010;14(1):20–35.

47. National Institute of Population Research and Training (NIPORT), International Centre for Diarrhoeal Disease Research, Bangladesh (icddr,b) and MEASURE Evaluation. Bangladesh Maternal Mortality and Health Care Survey 2016: Preliminary Report. Dhaka, Bangladesh and Chapel Hill, North Carolina, USA: NIPORT, icddr,b, and MEASURE Evaluation; 2017.

48. National Institute of Population Research and Training (NIPORT), Mitra and Associates, and ICF International. Bangladesh Demographic and Health Survey 2014. Dhaka, Bangladesh, and Rockville, Maryland, USA: Mitra and Associates, and ICF International; 2016.

49. Lesnewski R. Preventing unintended pregnancy: implications for physicians. American family physician. 2004;69(12):2779–80.

50. Adewuya AO, Fatoye FO, Ola BA, Ijaodola OR, Ibigbami S-MO. Sociodemographic and obstetric risk factors for postpartum depressive symptoms in Nigerian women. Journal of Psychiatric Practice®. 2005;11(5):353–8.

51. Adewuya AO, Ola BA, Aloba OO, Dada AO, Fasoto OO. Prevalence and correlates of depression in late pregnancy among Nigerian women. Depression and anxiety. 2007;24(1):15–21.

52. Giannandrea SA, Cerulli C, Anson E, Chaudron LH. Increased risk for postpartum psychiatric disorders among women with past pregnancy loss. Journal of Women’s Health. 2013;22(9):760–8.

53. Buist A. Perinatal depression--assessment and management. Australian family physician. 2006;35(9):670.

54. Britton JR. Maternal anxiety: course and antecedents during the early postpartum period. Depression and anxiety. 2008;25(9):793–800.

55. Schwab-Reese LM, Schafer EJ, Ashida S. Associations of social support and stress with postpartum maternal mental health symptoms: Main effects, moderation, and mediation. Women & health. 2017;57(6):723–40.

56. Li Y, Zeng Y, Zhu W, Cui Y, Li J. Path model of antenatal stress and depressive symptoms among Chinese primipara in late pregnancy. BMC pregnancy and childbirth. 2016;16(1):180.

57. Beydoun HA, Beydoun MA, Kaufman JS, Lo B, Zonderman AB. Intimate partner violence against adult women and its association with major depressive disorder, depressive symptoms and postpartum depression: a systematic review and meta-analysis. Social science & medicine. 2012;75(6):959–75.

58. Wu Q, Chen H-L, Xu X-J. Violence as a risk factor for postpartum depression in mothers: a meta-analysis. Archives of women’s mental health. 2012;15(2):107–14.

59. Howard LM, Oram S, Galley H, Trevillion K, Feder G. Domestic violence and perinatal mental disorders: a systematic review and meta-analysis. PLoS medicine. 2013;10(5):e1001452.

60. Ludermir AB, Lewis G, Valongueiro SA, de Araújo TVB, Araya R. Violence against women by their intimate partner during pregnancy and postnatal depression: a prospective cohort study. The Lancet. 2010;376(9744):903–10.

61. Bacchus L, Mezey G, Bewley S. Domestic violence: prevalence in pregnant women and associations with physical and psychological health. European Journal of Obstetrics and Gynecology and Reproductive Biology. 2004;113(1):6–11.

62. Kabir ZN, Nasreen H-E, Edhborg M. Intimate partner violence and its association with maternal depressive symptoms 6–8 months after childbirth in rural Bangladesh. Global health action. 2014;7(1):24725.

63. Ziaei S, Frith AL, Ekström E-C, Naved RT. Experiencing lifetime domestic violence: associations with mental health and stress among pregnant women in rural Bangladesh: the MINIMat Randomized Trial. PLoS one. 2016;11(12):e0168103.

64. Capaldi DM, Knoble NB, Shortt JW, Kim HK. A systematic review of risk factors for intimate partner violence. Partner abuse. 2012;3(2):231–80.

65. Axelson O, Fredriksson M, Ekberg K. Use of the prevalence ratio v the prevalence odds ratio as a measure of risk in cross sectional studies. Occupational and environmental medicine. 1994;51(8):574.

66. Lee J. Odds ratio or relative risk for cross-sectional data? International Journal of Epidemiology. 1994;23(1):201–3.

67. Angeles G, Lance P, Barden-O’Fallon J, Islam N, Mahbub A, Nazem NI. The 2005 census and mapping of slums in Bangladesh: design, select results and application. International Journal of Health Geographics. 2009;8(1):32.

68. Alam N, Begum D, Ahmed SM, Streatfield PK. MANOSHI COMMUNITY HEALTH SOLUTIONS IN BANGLADESH-Impact Evaluation Surveys in Dhaka Urban Slums 2007, 2009 and 2011. 2011.

